# Respiration and individual differences shape behavioral and inter-brain synchronization

**DOI:** 10.64898/2026.09.28.755199

**Authors:** Wenbo Yi, Caroline Palmer

## Abstract

Humans coordinate their actions while their bodies and brains synchronize in social interactions, yet how behavioral, physiological, and neural dynamics interact during interpersonal coordination remains largely unknown. We investigated how respiration and individual differences influence behavioral and inter-brain synchrony in a musical synchronization task, which requires precise temporal alignment. Eighty-eight individuals first produced a familiar melody at their most comfortable rate to measure their spontaneous production rate (SPR). They then synchronized the melody with a randomly assigned unfamiliar partner under natural (unpaced), instructed in-phase, and instructed anti-phase breathing conditions, while hyperscanning 64-channel electroencephalography (EEG), respiration, auditory-motor timing, and perceived social connectedness were recorded. Relative to natural respiration, both in-phase and anti-phase breathing increased inter-brain phase-locking and perceived social closeness but reduced behavioral synchrony; the instructed respiratory phase relationship was mirrored in their inter-brain relative phase. Differences in phase of respiratory cycles (inhalation and exhalation) also influenced coordination: Inhalation was associated with stronger inter-brain synchronization but less precise and stable behavioral coordination than was exhalation. Moreover, Individual differences influenced coordination: Partners with the faster SPR consistently preceded the slower partners and exhibited an alpha-band inter-brain phase advance in synchronization, whereas dyads with similar SPR showed greater behavioral synchrony and smaller inter-brain phase difference. Our findings identify respiration as a previously overlooked factor that organizes behavioral and neural synchronization, while individual intrinsic rhythms constrain how interacting partners align in time. Together, these findings reveal a dynamic interplay between the rhythms of the body and brain during human social interactions.

**Significance statement:** Respiration is a fundamental physiological process that shapes human cognition, yet its role in social interaction is unknown. We studied randomly paired dyads who synchronized music together, a human social behavior that requires fine temporal coordination. We found that respiratory rate, respiratory phase, and the inhale-exhale cycle shaped both behavioral and inter-brain dynamics. Moreover, people varied substantially in the natural rate at which they spontaneously produced music. Individual differences in this intrinsic frequency predicted both behavioral and inter-brain coordination in the dyads’ performances. These findings highlight the active role of the body-brain connection in human social interaction, and establish causal evidence disentangling behavioral, physiological, and neural processes in joint action through respiration.

## 1. Introduction

Interpersonal coordination, in which individuals temporally synchronize their actions, is a fundamental mechanism that facilitates human social interaction (1–5). Synchrony in coordination can occur across timescales and span behavioral, physiological, and neural processes (1, 6–11). Inter-brain synchrony (IBS), the temporal alignment of neural activity across individuals, is often considered a biological marker of successful interaction (12–16), with increased IBS being associated with more synchronised behaviour in joint action (7, 17, 18). However, recent findings suggest that the relationship between IBS and behavioral coordination may depend on the task and context. For example, greater behavioral instability in dyadic tapping tasks was associated with stronger theta-band IBS in central and left-temporal areas (19), whereas piano-duet performance with greater behavioral asynchrony and variability was associated with increased delta- and theta-band IBS (20). Such inconsistent relationships between inter-brain and behavioral synchrony may reflect other variables, such as physiological rhythms including respiration or heart rate, that can affect both behavior and brain but have been largely overlooked in interpersonal coordination.

Respiration is a plausible mechanism that may influence behavioral and inter-brain synchrony. Within individuals, respiration entrains neural oscillations across widespread brain networks (21, 22), and modulates cognitive and sensorimotor processes (23–29), which varies systematically with both respiratory frequency and phase. Inhalation often enhances perceptual and cognitive performance (30), whereas self-initiated motor actions tend to align with exhalation (28). Faster respiration rate shortens reaction and movement times, and a slow rate is associated with higher cognitive flexibility (31–33). Recent work showed that perturbing respiration during joint coordination impaired behavioral synchrony, supporting a causal influence of respiration on behavioral coordination (9). Whether respiration influences inter-brain interrelations is unknown. We address how the frequency and phase relationships of respiration between partners influence their behavioral and inter-brain coordination.

Individual differences in rhythmic behavior may also influence inter-brain synchrony. Individuals differ reliably in the spontaneous rates at which they produce auditory rhythmic sequences such as speaking, singing, and music performance (34–38). These rates, known as spontaneous production rates (SPRs), reflect intrinsic timing preferences that minimize energy expenditure and have been modeled as the natural frequencies of internal oscillators governing action timing (39–41). During musical coordination, partners with more similar SPRs synchronize more accurately, whereas the partner with the faster SPR tends to lead the slower partner behaviorally (9, 36, 42). Whether individual differences in intrinsic timing shape inter-brain dynamics remains largely unknown.

The current study tested whether respiration and individual differences influence dyads’ behavioral and inter-brain coordination during a musical synchronization task. Eighty-eight adults, randomly paired into 44 dyads, performed duets that required partners to precisely coordinate the timing of melody tones. Participants’ individual SPR were first measured in individual tasks; then partners performed the dyadic melody synchrony task while both partners’ neural activity (64-channel EEG), thoracic respiration, auditory-motor tone onsets, and perceived social connectedness ratings were recorded (Figure 1A). The relationship between the partners’ respiratory rhythms was experimentally manipulated across three conditions: Normal respiration, in which both partners were given no breathing instructions; In-phase respiration, in which partners inhaled and exhaled through their noses together at prescribed locations in the melody; and Anti-phase respiration, in which partners breathed at the same rate and melody locations as the In-phase condition while inhaling/exhaling in opposite phases (Figure 1B).

**Fig. 1.**
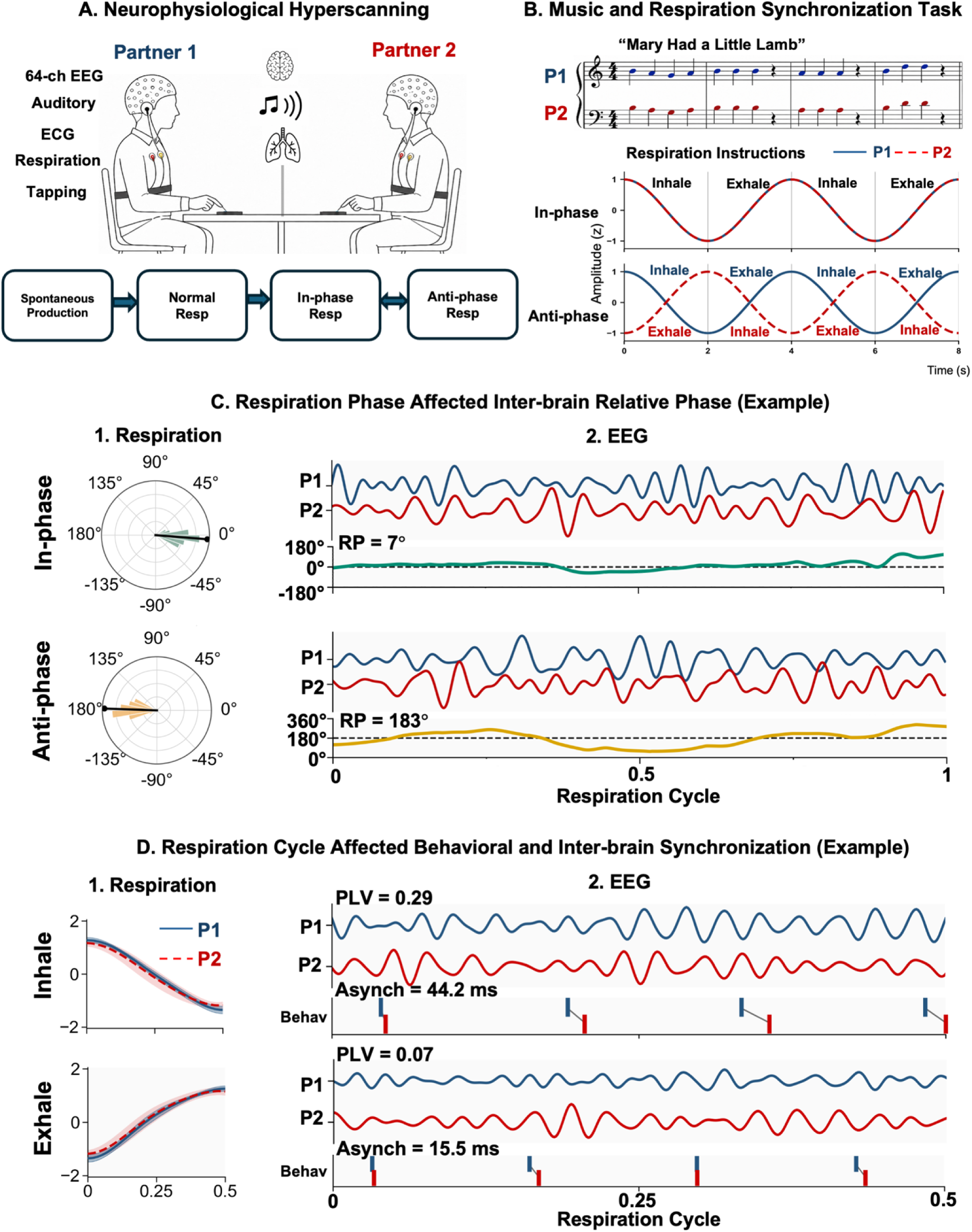
Experimental design and respiratory, behavioral, and inter-brain phase relationships during dyadic musical synchronization. (A) Schematic of the hyperscanning setup and experimental procedure. Partners synchronized melodic performances while EEG, ECG, respiration, and tapping were recorded simultaneously. Individual spontaneous production rate measurements preceded the Normal, In-phase, and Anti-phase respiration conditions (dyadic synchronization task); the order of the two instructed-respiration conditions was counterbalanced across dyads. (B) Musical task and schematic respiratory instructions. Partners performed “Mary Had a Little Lamb” together while inhaling and exhaling every four beats, either in phase or in opposite phases. Blue and red indicate partners 1 and 2, respectively. (C) Respiratory and inter-brain relative phase (RP; partner 1 minus partner 2). (1) Condition-level distributions of mean respiratory RP. Colored bars show the circular distribution of dyad-level respiration RP, and black vectors indicate the circular mean direction and concentration. (2) Example of theta-band EEG signals at homologous electrodes and corresponding inter-brain RP over one respiratory cycle from the same dyad in In-phase and Anti-phase conditions. RP was 7° in the In-phase example and 183° in the Anti-phase example. Dashed lines indicate 0° and 180° reference offsets. In the example, dyadic EEG relative phase mirrored the respiratory phase relationship. (D) Example of neural and behavioral synchronization during inhalation and exhalation cycles in the In-phase condition. (1) Mean z-scored respiration signals across dyads, with shaded areas indicating between-dyad SD. (2) Example alpha-band EEG signals at homologous electrodes and corresponding tapping events, selected from different respiratory cycles of the same trial and dyad shown in (2). Vertical marks indicate partners’ tap onsets, and connecting lines identify matched taps. In this trial, inter-brain phase-locking value (PLV) and mean absolute asynchrony were smaller (more synchronous) during exhalation compared to inhalation.

Two complementary metrics were applied to the dyads’ respiratory and inter-brain measures: Phase-locking values (PLV) quantified the consistency of phase differences between the partners’ signals (7, 9, 43–45), while mean relative phase (RP) measured the average phase offset and indicated the directionality of the partners’ signals (44, 46–48). We expected both in-phase and anti-phase breathing to increase inter-brain PLV relative to Normal breathing, as shared breathing rates may entrain partners’ neural oscillations (22, 23). We also expected In-phase and Anti-phase breathing to produce distinct inter-brain relative-phase patterns because respiratory phase can organize the phase of neural oscillations (24, 29). Behavioral (melodic tone) synchrony was expected to decrease under instructed breathing due to additional task demands. Partners with similar SPRs were expected to show greater behavioral and inter-brain synchrony, with faster-SPR partners leading slower partners in both behavioral timing and inter-brain relative phase.

## 2. Results

### 2.1 Respiration manipulation altered dyadic respiratory synchrony and rate

We first confirmed the In-phase and Anti-phase breathing manipulations with the respiration PLV and mean RP. Repeated-measures ANOVA revealed a significant respiration condition effect on the dyadic respiratory PLV, *F*(1.28, 55.18) = 633.32, *p* < .001, η² = .907 (G-G corrected). Pairwise comparisons showed higher PLV in both the In-phase and Anti-phase conditions than in the Normal condition (both *p_BH_* < .001); the In-phase and Anti-phase conditions did not differ (*p_BH_* = .531). Furthermore, a repeated-measures Hotelling’s test revealed a significant respiration effect on circular phase direction in dyadic RP, *T*² = 21792.49, *F*(4, 40) = 5,068.02, *p* < .001. Specifically, mean RP values were −10.44° in the Normal condition (90% CI, [−47.94°, 27.06°]), −4.52° in the In-phse condition (90% CI, [−8.63°, −0.42°]), and 177.99° in the Anti-phase condition (90% CI, [174.01°, 181.98°]). Finally, a repeated-measures ANOVA revealed a significant respiration effect on respiration rates (F(1.29, 55.29) = 8.92, *p* = .0021, η² = .083, G-G corrected). Pairwise comparisons showed lower respiration rates in both the In-phase (mean = 18.1 BPM, *p_BH_* = .007) and Anti-phase (mean = 18.3 BPM, *p_BH_* = .007) conditions than in the Normal condition (mean = 19.4 BPM), whereas the In-phase and Anti-phase conditions did not differ (*p_BH_ = .34*). Thus, relative to Normal breathing, both In-phase and Anti-phase respiratory manipulations increased dyadic respiratory phase synchrony during the intended in-phase and anti-phase relationships, and yielded slower breathing rates.

### 2.2 Respiration affected Auditory-motor synchrony and perceived social cohesion

We next examined whether respiration instructions altered the temporal precision of dyadic auditory–motor coordination or social cohesion ratings. The initial metronome cue of 500-ms intertone intervals was mirrored in all conditions: The partners’ mean intertone intervals (ITIs) were 480.90 ms (SE = 9.22 ms) in the Normal condition, 488.63 ms (SE = 9.19 ms) in the In-phase condition, and 487.81 ms (SE = 8.85 ms) in the Anti-phase condition. Mean ITIs did not differ significantly across conditions, F(1.35, 57.99) = 1.24, *p* = .284, η²= .003 (G–G corrected). Partners’ tone-onset absolute asynchronies (intended to be 0) and the SD of signed asynchronies were measured as indices of temporal accuracy and variability, respectively. Mean absolute asynchrony differed significantly across conditions (Figure 2A), F(1.68, 72.21) = 9.57, *p* < .001, η² = .044 (G–G corrected). Relative to the Normal condition, mean absolute asynchrony was higher in both the In-phase and Anti-phase conditions (*p_BH_* = .0034 and .0023, respectively), whereas the two instructed-respiration conditions did not differ (*p_BH_*= .758). Temporal variability showed the same pattern: the SD of signed asynchrony differed across conditions, F(2, 86) = 6.64, *p* = .0021, η²= .068, with higher values in the In-phase and Anti-phase conditions than in Normal (*p_BH_* = .0101 and .0049, respectively), but no difference between In-phase and Anti-phase (*p_BH_* = .609). Thus, the instructed respiration conditions decreased behavioral synchronization relative to uninstructed (natural) respiration.

**Fig. 2.**
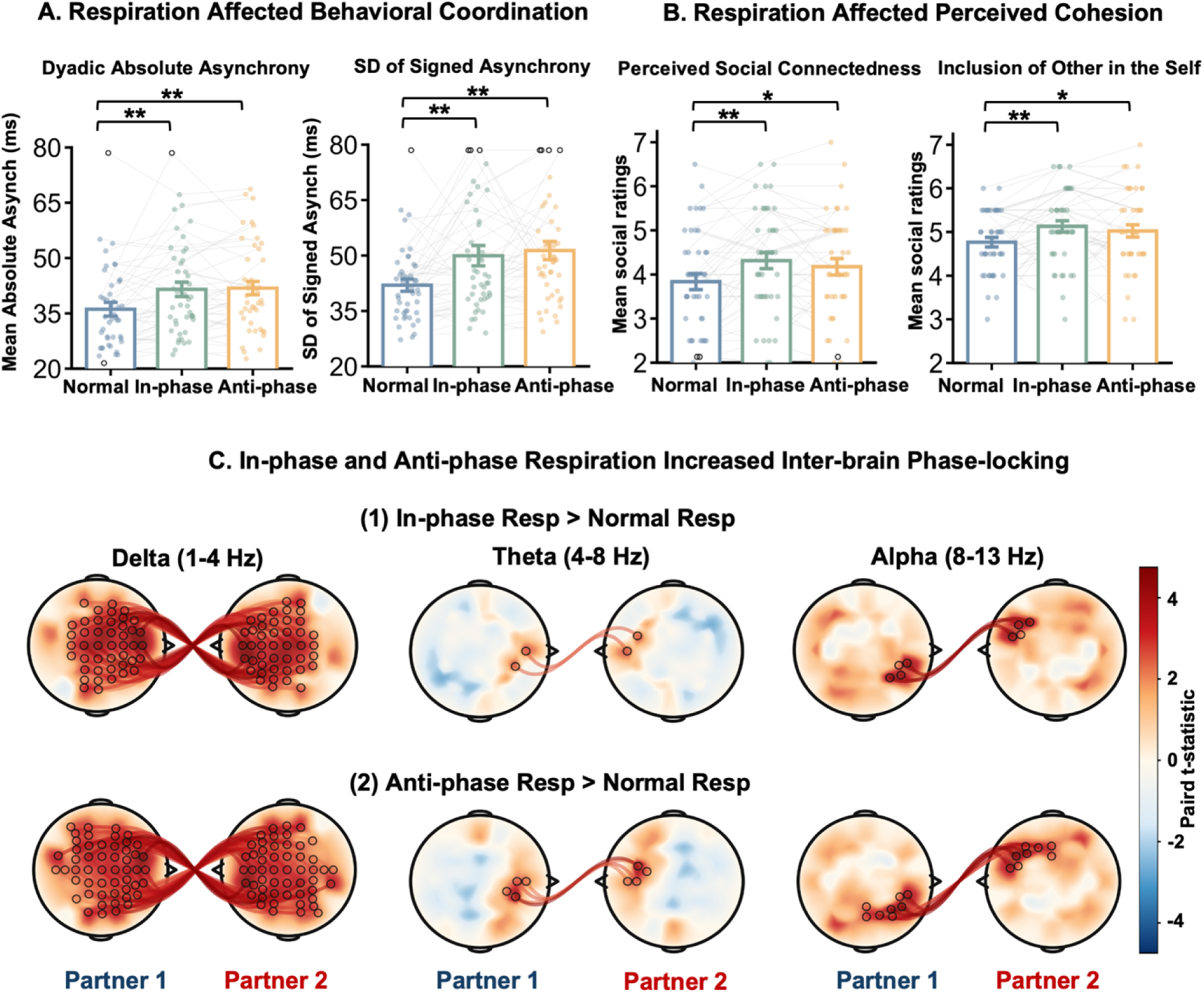
Instructed respiration reduced behavioral synchrony while increasing perceived social connectedness and inter-brain phase locking. (A) Condition-level dyadic mean absolute asynchrony and SD of signed asynchrony in tone onsets. Both measures increased during In-phase and Anti-phase respiration relative to Normal breathing, indicating less accurate and less stable auditory–motor synchronization. (B) Condition-level perceived social connectedness and Inclusion of Other in the Self individual ratings. Both ratings increased during In-phase and Anti-phase respiration relative to Normal breathing. In A and B, bars show means ± SEM. Asterisks indicate Benjamini–Hochberg-corrected pairwise significance: *: *p* < .05 and **: *p* < .01. (C) Condition-level contrasts (cluster permutation test) in inter-brain phase-locking value (PLV) between homologous electrodes. In-phase (1) and Anti-phase (2) respiration increased PLV relative to Normal breathing in significant clusters spanning delta, theta, and alpha bands (all cluster-level *p’s* < .001). Colors indicate paired t-statistics; outlined electrodes and connecting lines identify homologous electrode pairs belonging to significant clusters.

Perceived social connectedness (“Overall, I feel connected to the person in the experiment with me”, with higher scores indicating a closer relationship) also differed between conditions (Figure 2B), Friedman χ²(2) = 10.99, *p* = .004. Conover–Friedman post hoc tests showed higher ratings in the In-phase (*p_BH_* = .0025) and Anti-phase (*p_BH_*= .045) conditions than in Normal breathing, whereas In-phase and Anti-phase did not differ (*p_BH_* = .211). Inclusion of Other in the Self (IOS) ratings (higher scores indicate a closer perceived social relationship) also differed across conditions, Friedman χ²(2) = 9.96, *p* = .0069. IOS ratings were higher in the In-phase (*p_BH_*= .005) and Anti-phase (*p_BH_* = .038) conditions than in Normal breathing, with no difference between In-phase and Anti-phase (*p_BH_*= .345). The remaining two social questions showed no significant condition effects (see SI.1). Thus, the instructed respiration conditions increased perceived social connectedness relative to natural respiration.

### 2.3 Respiration frequency affected inter-brain phase-locking

We tested whether the partners’ respiratory frequency altered inter-brain synchrony, quantified as PLV between homologous EEG electrode pairs. For each of the pairwise within-condition contrasts, paired t-statistics were evaluated using a cluster-based permutation method (7, 49–52), with 5,000 permutations across all 64 electrode pairs and four frequency bands (Delta, Theta, Alpha, and Beta) to minimize spatial selection bias and control the Family-Wise Error Rate (FWER, details see SI. 2). Relative to the Normal respiration condition, inter-brain PLV measures were higher in the In-phase respiration condition in one significant cluster (*p_FWER_* < .001), and were also higher in the Anti-phase condition (*p_FWER_*< .001, Figure 2C). Both in-phase and anti-phase respiration increased the inter-brain PLV in delta, theta, and alpha bands, and predominantly at lower frequencies relative to Normal breathing. There was no significant difference between the two instructed-respiration conditions (In-phase and Anti-phase, *p_FWER_* = .99), and no significant effect in the beta band. Therefore, synchronizing respiration frequency significantly increased inter-brain synchrony. Chance-level inter-brain phase coupling was assessed using both pseudo-dyad and within-trial epoch surrogates (49). Observed PLV exceeded surrogate estimates in all three conditions (all *p* < .001), indicating the observed PLV reflects real dyadic interaction and above-chance inter-brain synchrony (see SI.3 for details).

### 2.4 Respiration phase relationship affected inter-brain relative phase

We examined whether the partners’ respiration relative phase altered inter-brain phase (RP) measures, quantified as the circular RP shift between homologous EEG electrode pairs. We contrasted the dyadic mean RP between the Anti-phase and In-phase respiratory conditions using a paired Hotelling’s T² test with the same 5,000-permutation cluster-based correction for multiple comparisons (Figure 3A). The contrast yielded one significant cluster (peak T² = 12.99 at theta/CP1, *p_FRER_*= .014). The effect was strongest in the theta band over the centroparietal region, with the largest T² values at CP1 (mean RP = 174.0°). The alpha band showed the broadest spatial involvement across central and centroparietal regions, peaking at C1 (T² = 12.60; mean RP = −98.8°), whereas the beta band was more focal over frontocentral and central regions, with a peak at C1 (T² = 11.28; mean RP = 166.2°). These findings indicate that the dyad’s respiratory relationship altered the spatial organization and direction of inter-brain phase relationship across frequency bands. The near-180° condition-related shifts in theta and beta suggest a pronounced reorientation of dyadic neural phase relationships affected by the partners’ respiration (In-phase and Anti-phase).

**Fig. 3.**
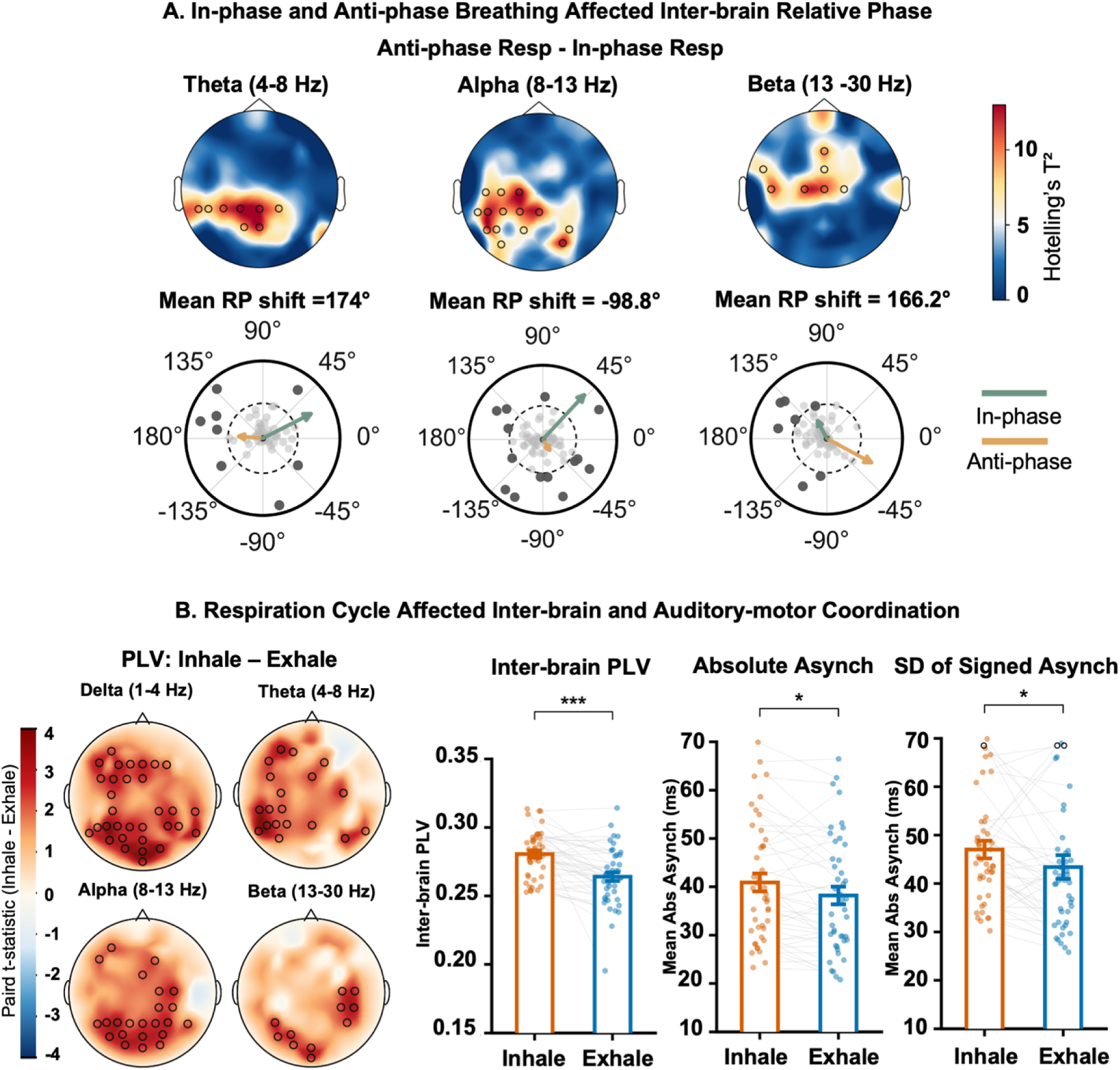
Dyadic respiratory coordination shaped inter-brain relative phase, and joint inhalation was associated with stronger neural synchrony and weaker behavioral synchrony. (A) Condition-level differences in inter-brain relative phase (RP) between Anti-phase and In-phase respiration across dyads. Topographies show paired Hotelling’s T² statistics, with outlined electrodes belonging to a significant cluster spanning theta, alpha, and beta bands (*p* = .014). Mean condition-related RP shifts across theta, alpha, and beta bands at significant electrodes were 174°, −98.8°, and 166.2°, respectively. In the polar coordinate graphs, each point represents a homologous electrode across partners: angle indicates the condition-related RP shift between partners, radial distance indicates T², and darker points identify significant cluster members identified by the cluster permutation test. Black dashed circles mark the cluster-forming threshold; colored arrows summarize condition-level mean partners’ phase vectors across significant electrodes. (B) Inhale–exhale comparisons in the In-phase respiration condition across dyads. Topographies show greater inter-brain phase-locking value (PLV) during inhalation in a significant cluster spanning all four frequency bands (*p* < .001); outlined electrodes identify cluster members. Bar charts summarize mean PLV values across the significant band–electrodes in this cluster. Mean absolute asynchrony and SD of signed asynchrony were smaller during exhalation, indicating more accurate and stable auditory–motor synchronization. Bars show means ± SEM and dots represent dyads. *: *p* < .05 and *** *p* < .001.

### 2.5 Within-cycle respiration affected inter-brain and auditory-motor synchronization

We then investigated whether different phases in the respiratory cycle affected inter-brain synchrony; we first compared inhalation and exhalation periods (when the partners shared the same respiratory cycle) in the In-phase and Normal conditions. Because PLV is sensitive to window size (52), we extracted and resampled each complete inhale-exhale period into a common normalized time, spanning one respiratory half-cycle for each inhale and each exhale period (53, 54). PLV was estimated from the normalized respiratory cycles and compared between homologous electrode pairs using the same cluster-based permutation test. Inter-brain PLV was higher during inhalation than exhalation in one significant cluster across all four frequency bands (*p_FWER_* < .001). To examine whether this pattern was present during both instructed and spontaneous respiratory coordination, we next analyzed the In-phase and Normal conditions separately. Both conditions showed higher PLV during inhalation than exhalation (In-phase: *p_FWER_* < .001; Normal: *p_FWER_*<.001; Figure 3B). Cluster-member electrodes were distributed predominantly over parietal and occipital regions in the In-phase condition, whereas the Normal cluster extended across all 64 electrodes in all four frequency bands. To assess whether this pattern was specific to partners’ synchronized respiration phase instead of the respiratory instruction, we applied the same pipeline to the Anti-phase condition, where partners are breathing in opposite phases. No significant cluster was detected in this condition (minimum *p_FWER_* = .120). These results indicate that dyads’ synchronized respiration was associated with greater inter-brain PLV during inhalation than during exhalation.

We next examined whether inhalation and exhalation differences were associated with dyadic auditory–motor synchronization. We extracted melody tone events that were produced in the partners’ same shared inhalation and exhalation periods. Because the mean absolute asynchronies and SD of signed asynchrony deviated substantially from normality (Shapiro-Wilk test, *p* < .05), non-parametric Wilcoxon signed-rank tests were applied. Both mean absolute asynchrony (*W* = 196, *p* < .001) and the SD of signed asynchrony (*W* = 229, *p* = .0015) were greater during inhalation than exhalation. Similarly, we next analyzed the In-phase and Normal conditions separately. Both conditions showed greater absolute asynchrony and variability during inhalation than exhalation (In-phase: absolute asynchrony, *W* = 279, *p* = .010; SD, *W* = 282, *p* = .012 (Fig 3B); Normal: absolute asynchrony, *W* = 245, *p* = .003; SD, *W* = 192, *p* < .001). Neither measure differed significantly in the Anti-phase condition (absolute asynchrony: *W* = 426, *p* = .428; SD of signed asynchrony: *W* = 428, *p* = .441). To further confirm whether the variability of the respiratory cycle changed, we calculated the SD of respiratory amplitude within each inhale and exhale cycle. Respiratory amplitude SD was significantly greater during inhalation than during exhalation in both Normal (W = 0, *p* < .001) and In-phase conditions (W = 181, *p* < .001). Together, these results indicate that inhalation cycles yielded greater variability in respiratory amplitude and more variable behavioral asynchrony than exhalation cycles in both Normal and In-phase respiration conditions.

### 2.6 Individual differences predict auditory-motor and inter-brain coordination

Finally, we assessed whether individual differences in auditory-motor production, measured by spontaneous production rates (SPR), influenced interpersonal coordination. Individual SPR values (Mean ITI) varied substantially (Fig 4A), ranging from 243.98 to 637.95 ms with a mean within-participant SE of 3.3 ms. To test whether the faster-SPR partner showed a directional relationship to the slower-SPR partner in auditory-motor synchronization, we compared their mean signed asynchrony (faster-SPR partner − slower-SPR partner) against zero using one-sample t-tests. Mean signed asynchronies in each condition were significantly negative (Figure 4B, Normal: t(43) = −2.92, *p* = .0056; In-phase: t(43) = −3.32, *p* = .0018; Anti-phase: t(43) = −2.65, *p* = .0112), indicating that faster-SPR participants consistently tapped ahead of the slower partner in the dyadic synchronization task. In addition, the signed SPR difference was positively associated with the auditory–motor asynchrony values (Figure 4C) in the Normal (Pearson *r* = 0.318, *p* = .0357), In-phase (Pearson *r* = 0.374, *p* = .0123), and Anti-phase (Pearson *r* = 0.478, *p* = .0010) conditions, indicating that dyads with similar SPRs achieved better synchrony in the dyadic synchronization task.

**Fig. 4.**
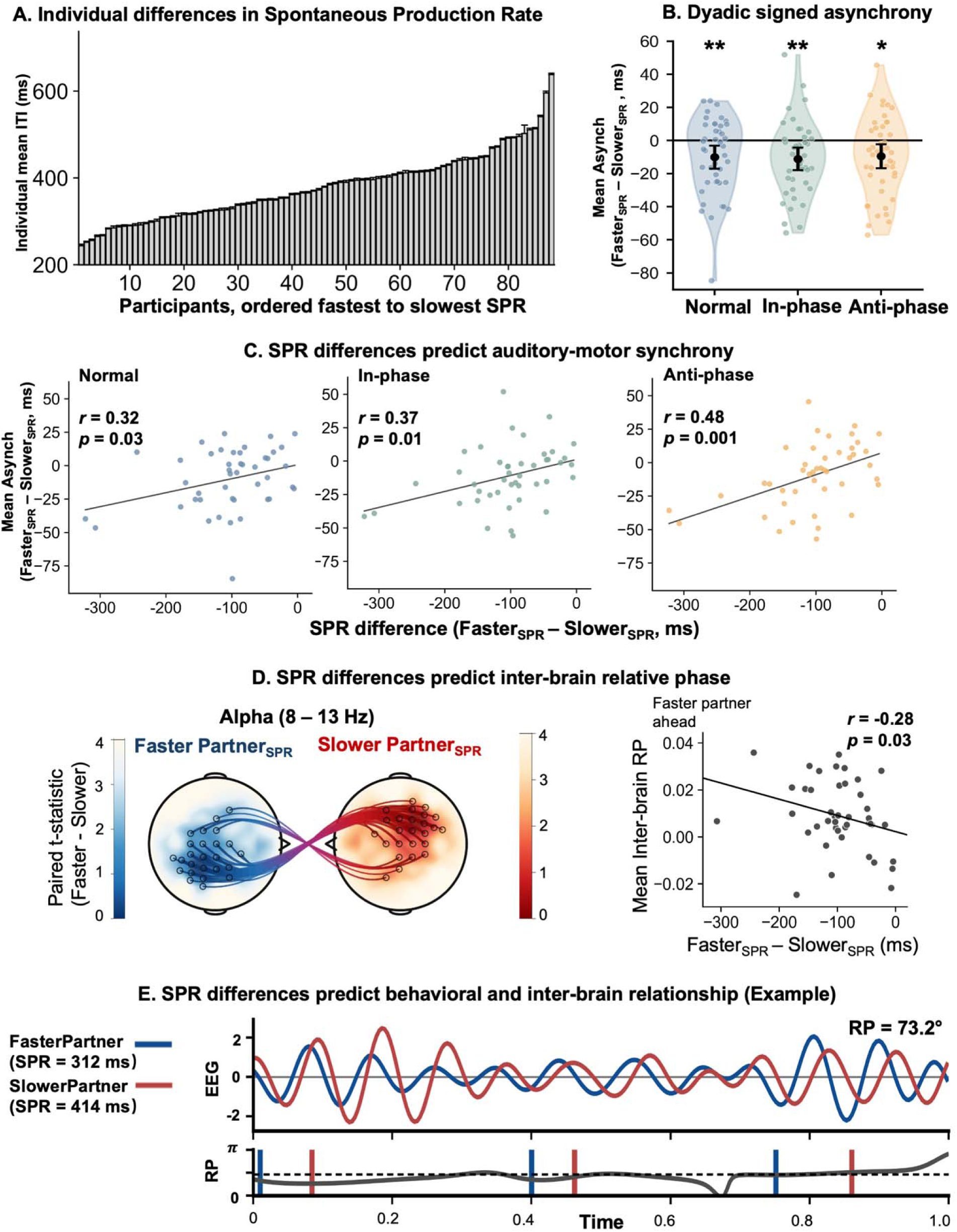
Individual differences in spontaneous production rate predicted behavioral and inter-brain phase relationships. (A) Spontaneous production rate (SPR), measured as mean intertap interval (ITI) during individual melody production (first task in study) by participant, ordered from smallest to largest ITI (fastest to slowest). Bars represent individual means, with error bars indicating SE of means. (B) Condition-level mean signed asynchrony across dyads, defined as the faster-SPR partner’s tone onset minus the slower-SPR partner’s tone onset. Negative values indicate that the faster-SPR partner led behaviorally. Violin plots show dyad-level distributions; black points and error bars indicate means and 95% confidence intervals. Faster-SPR partners preceded across all three Respiration conditions. Asterisks indicate two-sided one-sample t-tests against zero: *: *p* < .05 and **: *p* < .01. (C) Individual SPR differences (A) predicted dyads’ signed asynchrony (B), with larger dyadic SPR differences predicting greater behavioral leading by the faster-SPR partner. Points represent dyads; lines show condition-level linear fits of Pearson correlations with p-values. (D) SPR differences predicted dyads’ inter-brain phase leading during the Normal condition. Condition-level mean relative phase values; Faster-SPR partners showed an alpha-band phase lead in a significant cluster of 25 homologous electrode pairs (*p* = .012). Outlined electrodes and connecting lines identify cluster members; red and blue distinguish faster- and slower-SPR partners, with color intensity indicating t-statistics. The scatterplot shows SPR differences predicted the cluster mean inter-brain relative phase of the faster-minus-slower Partner’s EEG. Positive y-axis values indicate faster-partner phase advance. (E) Example dyadic trial (one-second segment) from the Normal condition showing z-scored alpha-band EEG at CP4 in both partners and the corresponding relative phase. Red and blue traces denote the faster- and slower-SPR partners, respectively; vertical lines mark the tapping events. The faster-SPR partner showed a mean inter-brain phase lead of 73.2° during this segment.

We next examined whether individual SPR differences influenced partners’ inter-brain RP values during synchronization. We calculated the sine of the EEG RP (faster-SPR partner − slower-SPR partner). Positive values of the signed relative phase indicated a lead in synchronization by the faster SPR partner. The same cluster-permutation test with one-sample t-tests was applied to assess whether the signed values were greater than zero. A significant positive alpha-band cluster in the central-right parietal region, peaking at CP4 and comprising 25 electrodes (*p_FWER_* = .0116), was identified, indicating that the faster-SPR partner had a significant cortical phase advance relative to the slower-SPR partner in the dyadic task (Figure 4C). Similarly, we then correlated the dyad-level SPR difference (faster partner ITI – slower partner ITI, ms) with the mean sine of the faster-minus-slower RP across the significant cluster. There was a significant negative correlation (r = −.285, *p* = .0322, one-sided), indicating that larger faster–slower SPR differences were associated with larger brain phase advance by the SPR-faster participant (Figure 4D). Thus, individual differences in auditory-motor production were associated with both the direction and strength of interpersonal coordination: faster-SPR partners led behaviorally and showed an alpha-band neural phase lead in synchronization, whereas dyads with similar SPR showed greater behavioral synchrony and smaller inter-brain RP differences.

## 3. Discussion

Human interpersonal synchronization emerges across behavioral, physiological, and neural levels. Previous studies have largely overlooked the roles of respiration and behavioral individual differences in interpersonal coordination. By experimentally manipulating dyadic respiration during musical synchronization, we showed that respiration influenced their perceived social connectedness, and the strength and direction of their inter-brain synchrony. Both in-phase and anti-phase respiration instructions increased inter-brain PLV relative to natural breathing, while cortical measures of relative phase (RP) mirrored the instructed respiratory phase relationships across partners. Inhalation–exhalation cycles also influenced both behavior and brain: inhalation was associated with higher inter-brain synchronization but less accurate and less stable behavioral synchrony. Spontaneous production rate (SPR), an individual measure of natural frequencies in auditory-motor production, also predicted behavioral and inter-brain synchrony, with faster partners leading slower partners in both auditory-motor behavior and inter-brain phase advances. Together, these findings show that inter-brain dynamics track physiological and behavioral dynamics, revealing how respiration and individual intrinsic rhythms shape human social behavior.

Perceived social cohesion and interbrain synchrony were enhanced in the In-phase and Anti-phase conditions relative to Normal (uninstructed) breathing, even though the respiration instructions increased task demands and reduced behavioral tone synchronization. At the cortical level, both respiration instructions produced widespread bilateral increases in inter-brain PLV, consistent with single-brain evidence that respiration modulates oscillatory activity in limbic and distributed cortical networks (22, 29). Stronger inter-brain synchrony thus coexisted with less accurate behavioral coordination. The dissociation may reflect increased task demands: Breathing instructions in addition to tone onset synchronization may have forced dyads to plan multiple movements (fingers and lungs). This interpretation is also supported by decreased behavioral synchrony during the inhale portion of respiratory cycles (which indicated that respiration planning did influence behavioral synchrony). Together, these findings suggest that synchrony in dyadic interaction is not a unitary phenomenon (10, 55); different levels of synchrony may index complementary facets of human interaction rather than a single shared temporal process.

The inter-respiratory phase relationship also shaped inter-brain relative phase across frequency bands and scalp regions. This organization is consistent with evidence that respiratory-brain phase modulation varies across frequency bands and brain regions, including a pronounced parieto-occipital effect (21, 22, 26). Although partners synchronized their actions to the same melody in both conditions, theta- and beta-band inter-brain relative phase still mirrored their contrasting respiratory phase relationships. This suggests that respiratory coordination organizes the temporal structure of inter-brain dynamics even in the presence of the same external stimuli. Our findings therefore extend evidence for individual respiration–brain coupling to dyadic interaction and identify coordinated physiology as an active, causal, and previously overlooked dimension of social interaction.

Differences in respiratory inhalation and exhalation phases also affected interpersonal coordination. Joint inhalation was associated with stronger inter-brain phase-locking but less precise and stable auditory-motor synchronization than exhalation. The heightened inter-brain synchrony may reflect increased cortical excitability across sensory and motor networks during inhalation (23, 26), potentially enhancing neural readiness and sensitivity to one another’s timing cues and supporting mutual adjustment. This pattern may also suggest increased cognitive effort required to plan and coordinate respiration, as reflected in greater respiratory amplitude variability. This interpretation is supported by evidence that reaction times to auditory stimuli were longer during inhalation than during exhalation (56), as well as by previous findings that greater coordination demands can increase inter-brain synchrony even when behavioral synchronization is perturbed (20). Exhalation, on the other hand, was associated with more precise and stable behavioral synchrony, broadly consistent with evidence that self-initiated motor actions occur preferentially during exhalation (28). These findings may suggest that inhalation supports preparation and ongoing timing adjustment, whereas exhalation favors more stable execution in coordination. Thus, our results demonstrate that the respiratory cycle affects neural and behavioral coordination differently and dynamically organizes auditory-motor preparation, adjustment, and execution during synchronization.

Individual differences in the spontaneous rates at which participants produced the same melody separately (at the beginning of the study) shaped partners’ behavioral and inter-brain coordination. Partners with faster SPRs consistently led the joint action, and dyads with more similar SPRs achieved more accurate auditory–motor synchronization. The asymmetry between faster and slower partners was also evident at the cortical level: Faster-SPR partners showed an alpha-band phase lead, with a topography near the right temporoparietal junction (rTPJ), a region implicated in integrating behavioral timing and social information in inter-brain relationships (57), than slower-SPR partners. The findings are in alignment with the neural resonance theory, which conceptualizes temporal attention as an internal oscillator that entrains to external rhythms (40, 58–60). Given the established role of alpha-band oscillations in attention (61–63), the faster partner’s cortical oscillation may reflect a phase-advanced attentional oscillator.

In sum, dyadic interpersonal coordination measured across behavioral, physiological, and neural levels revealed the active role of respiration and behavioral intrinsic production frequencies in engaging distinct cortical frequency bands. Future work may combine EEG with slower hemodynamic imaging to link rapid inter-brain dynamics with the slower changes present in respiration cycles, to address how interpersonal coordination emerges, stabilizes, and adapts over time.

## 4. Methods

### 4.1 Participants

Eighty-eight healthy adults aged 18–34 years (M = 20.0, SD = 2.01; 7 males) were recruited from the Montreal community and randomly assigned to 44 dyads. The target sample of 44 dyads was determined a priori using G*Power, assuming a medium effect size (f = 0.25; partial η² = 0.06) and 90% statistical power.

All participants were strangers to each other and had at least two years of group or individual music training on a single instrument (years of musical training: M = 6.18 years, SD = 3.39). This inclusion criterion ensured the sensorimotor precision required to maintain synchronization while following the respiration instructions (9). Before the experiment, all participants passed an auditory perceptual screening confirming that they could detect tone-onset asynchronies for two tones differing in pitch similar to the melodies they would produce (SI. 4). Participants provided informed consent before beginning the experiment. The study protocol was approved by the McGill University Research Ethics Board. Participants received either course credit or a small honorarium for completing the experiment, which lasted approximately 2 hours.

### 4.2 Stimuli and Equipment

The auditory sequences consisted of a familiar melody, “Mary Had a Little Lamb,” sounded in two different octaves in the dyadic synchronization task (one assigned to each partner, beginning on B3 and B4). This octave separation allowed the two sequences to remain discriminable when sounded simultaneously in the synchronization task. The same melody was used in the spontaneous rate task. The melodies were presented with a classic piano timbre on a Roland Studio Canvas SD-50 tone generator. To ensure precise melody production timing, participants performed on Arduino-based force-sensitive tapping pads connected to a Linux computer via MIDI (64, 65). Each tap triggered the next tone in the stimulus melody, thus allowing participants to control the melody rate, and the tones were delivered through ER-1 insert earphones. Tap timing was recorded at 1-ms resolution using FTAP (9, 66).

Brain activity was recorded from each participant with a 64-channel BioSemi Active Two EEG system (BioSemi, Amsterdam, the Netherlands) synchronized via an optical daisy-chain configuration at 2048 Hz (67) in an electromagnetically shielded, sound-proofed EEG recording room (ETS-Lindgren, Texas, USA). Two additional electrodes were placed over the left and right mastoids for offline re-referencing. Electrode positions followed the international 10–20 system. Electrode offsets relative to the common mode sense (CMS) and driven right leg (DRL) feedback loop were maintained below ±40 mV (68, 69). Respiration was recorded using a force-sensitive respiration belt connected to the EEG AUX channel and secured around the chest near the diaphragm, providing a measure of relative respiratory effort. Cardiac activity was recorded as a bipolar ECG using two flat electrodes connected to external BioSemi channels and placed on the right waist and left upper chest (Cardiac synchrony results reported in SI. 5). Social interaction was assessed using four questions adapted from previous studies (9, 70), measuring perceived social connectedness, pleasantness, inclusion of the other in the self (IOS), and perceived success. Participants completed the ratings on paper using 7-point Likert scales, with higher scores indicating greater social cohesion (See SI.1).

### 4.3 Experimental Design and Procedure

The experiment followed a within-subject design in the following order: individual spontaneous production (SPR) task and a dyadic synchronization task with three conditions. The Normal (uninstructed) condition was completed first; the In-phase and Anti-phase condition orders were counterbalanced across dyads.

The procedure began with participants providing consent and completing a musical background questionnaire (9). Next, participants individually (separate from their partner) completed the standardized SPR task to determine the rate of their spontaneous production of a familiar melody, “Mary Had a Little Lamb” (36, 38, 42). All participants confirmed their familiarity with the melody. The mean and standard error of the intertap intervals (ITIs) were used to obtain each participant’s SPR (See SI.6 for SPR procedure and analysis).

Participants then completed the dyadic synchronization tasks together, with the melody notation in front of them. They produced the melody while seated face-to-face, with a visual barrier occluding their partners’ hand and arm movements below the chest (Fig. 1A). Participants were instructed to synchronize their melody with their partner as accurately as possible after an eight-beat auditory cue ended (500 ms ITI), while maintaining the cued rate. Practice trials preceded each condition. In the Normal condition, participants synchronized without respiratory instructions. In the In-phase condition, both partners synchronized with the same melody, but began using their nose to inhale on the first beat and exhale four beats later, alternating every four beats (one musical bar). In the Anti-phase condition, partners followed the same respiratory rate in opposing phases: one began inhaling while the other began exhaling. The partner assigned to begin with inhalation was counterbalanced across trials. After each respiration condition, participants completed the Social Interaction Questionnaire.

### 4.4 Data Analyses

#### 4.4.1 Neurophysiological Data Preprocessing

The neurophysiological data were extracted and processed using the MNE Python toolbox (71, 72) and were visually inspected by the authors for data quality. In the two respiratory manipulation conditions, respiration inhalations and exhalations were examined for compliance with the instructed In-phase and Anti-phase breathing patterns. Noncompliant periods were annotated for exclusion. Channel-specific EEG artifacts identified during inspection were repaired using spherical-spline interpolation, whereas intervals affecting multiple electrodes were rejected. Within dyads, rejection intervals identified in either participant were applied to both partners and across all modalities of data. The interpolation affected 0.416% of the total pre-rejection duration, and rejection removed 6.871% of the total trial durations in the final dataset.

Hyperscanning EEG signals were processed using procedures adapted from the HyPyP pipeline (49). Continuous signals were bandpass-filtered between 0.5 and 40 Hz, re-referenced to the averaged mastoids, and resampled to 256 Hz. Trials were cropped between the earliest and latest tapping markers. Then, the visually inspected signals were processed by independent component analysis (ICA), through which eye blinks and movements, muscular, and cardiac components were identified and removed. Cleaned continuous signals were divided into non-overlapping 2-s epochs, and further visually inspected to confirm the artifacts were being removed. The respiration data pre-processing followed established pipeline using Neurokit 2 (73) (See SI.7 for details).

#### 4.4.2 Inter-brain and respiratory synchrony analyses

To quantify inter-brain and respiratory synchrony between dyad members, two phase-based measures were applied to the EEG and respiratory signals. Phase-locking value (PLV), a widely used measure of phase-relationship consistency in EEG and respiratory analyses (7, 9, 43–45), was defined as

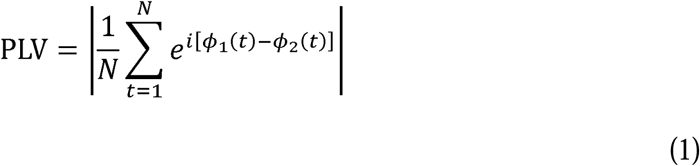

PLV quantified the stability of the dyadic relative phase within each modality between the dyadic members, ranging from 0 (dispersed phase differences) to 1 (a constant phase difference).

Mean relative phase (RP) was quantified to identify the direction of dyadic neurophysiological data (44, 46–48). For participant *p*, the instantaneous phase at sample *t* was denoted by *ϕp*(*t*) The dyadic relative phase was defined as partners’ phase difference (either SPR-defined faster partner – SPR-defined slower partner in SPR-related analysis; or partner 1, high-pitched partner – partner 2, low-pitched partner) and wrapped to the interval [-*π*, *π*):

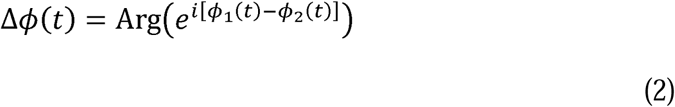

Values near 0 indicated an in-phase relationship, whereas values near ±*π* indicated an antiphase relationship.

EEG analyses were restricted to the delta (1–4 Hz), theta (4–8 Hz), alpha (8–13 Hz), and beta (13–30 Hz) bands. Gamma activity was excluded to reduce potential contamination from respiration-related cranial muscle activity (74, 75). Instantaneous phase was obtained using the Hilbert transform; mean relative phase and PLV were calculated between homologous electrodes (e.g., Cz–Cz and C2–C2) within each epoch, and then averaged across valid epochs within each trial. We focused on homologous electrode pairings across the dyad members to assess simultaneous phase alignment at corresponding scalp locations, which may capture shared distributed neural processes across dyad members (76).

#### 4.4.3 Behavioral data analysis

Tap-triggered tone onsets and intertap intervals (ITIs) were extracted from each partner’s force-sensor signals using custom Python scripts. Absolute asynchrony was calculated as the absolute difference between partners’ tap onsets at matched serial positions in the melody and averaged within each trial. The participant with the smaller mean ITI in the individual SPR task was classified as the faster partner. Signed asynchrony was calculated as the faster partner’s tap onset minus the slower partner’s tap onset; negative values indicated that the faster participant preceded the slower participant.

#### 4.4.4 Statistical Analysis

Behavioral and respiratory outcomes were analyzed at the dyad level using separate one-way repeated-measures analyses of variance (ANOVAs), with respiration condition (Normal, In-phase, and Anti-phase) as the within-dyad factor and α = .05. Sphericity was assessed using Mauchly’s test; when violated, Greenhouse–Geisser-corrected degrees of freedom and p-values were reported. Multiple comparisons were corrected by Benjamini–Hochberg (FDR) correction. Because social-connectedness responses were measured on a 7-point ordinal scale, respiration condition effects on each questionnaire item were assessed with Friedman tests, followed by Conover–Friedman pairwise comparisons. Correlations were assessed using Pearson’s correlation coefficient. Data points falling more than three SD from the mean were excluded as outliers. Across all correlation analyses, only one data point met this criterion and was excluded from the analysis examining the association between phase and SPR difference. The statistical significance of condition differences in EEG PLV and RP was determined using cluster-based permutation tests (7, 49, 51, 52), with family-wise error controlled across sensors and frequency bands (see SI.2 for details).

## Acknowledgement

The authors acknowledge the assistance of Liam Curry, Emilia Zarb, and Kai Mikkelsen in technical setup and data collection. This work was partially funded by grants from the Fonds de recherche du Québec Nature et technologies (FRQNT) Doctoral Fellowship to W.Y., and from the Natural Sciences and Engineering Research Council of Canada (Discovery Grant 298173) to C.P.

## Data Availability Statement

The de-identified data and code supporting the manuscript will be made available upon request during the peer-review process, and will be made fully available upon publication of the manuscript.

