## Supplementary material for "Respiration and individual differences shape behavioral and inter-brain synchronization": SI

**Supplementary Information**

Wenbo Yi^1,2^ & Caroline Palmer^1,2^

1. Department of Psychology, McGill University, QC, CA
2. Centre for Research on Brain, Language and Music, QC, CA

**SI.1 Social interaction questionnaires**

**The questions below concern your impression of the experiment you just completed. Please answer all questions by circling a number on each of the scales. There are no right or wrong answers to any of these questions, we are just interested in your opinions.**

1. How unpleasant or pleasant did you find the experience of playing with the other participant?

|  | 1 | 2 | 3 | | 4 | 5 | 6 | | 7 |
| --- | --- | --- | --- | --- | --- | --- | --- | --- | --- |
| Extremely unpleasant | | | |  | | | | Extremely pleasant | |

2. Overall, I feel connected to the person in the experiment with me.

|  | 1 | 2 | 3 | | 4 | 5 | 6 | | 7 |
| --- | --- | --- | --- | --- | --- | --- | --- | --- | --- |
| Disagree Strongly | | | |  | | | | Agree Strongly | |

3. Please circle the picture below that best describes how your relationship with the other person in the experiment currently is:

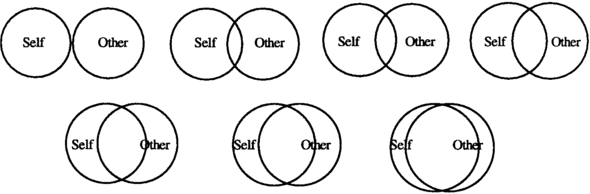

4. How successful do you think your synchronization was with your partner?

|  | 1 | 2 | 3 | | 4 | 5 | 6 | | 7 |
| --- | --- | --- | --- | --- | --- | --- | --- | --- | --- |
| Not at all  synchronized | | | |  | | | | Very well synchronized | |

Questions 2 (perceived social connectedness) and 3 (IOS ratings) showed significant effects, as reported in the main text, where as Questions 1 (experience pleasantness) and 4 (perceived synchronization success) showed no significant condition effects: Ratings of experience pleasantness (Q1: “How unpleasant or pleasant did you find the experience”) did not differ significantly across the Normal, In-phase, and Anti-phase conditions, Friedman χ²(2) = 3.45, p = .178. Ratings of perceived synchronization success (Q4) also showed no significant condition effect, Friedman χ²(2) = 2.03, p = .363. Both analyses used ratings averaged within each of the 43 dyads with complete questionnaire data (one pair of data is missing). Neither omnibus test was significant, so no post hoc comparisons were performed.

**SI.2 Inter-brain cluster-permutation analysis description**

Dyads' phase-locking values were analyzed on all 64 homologous sensor at four frequency bands with cluster-based permutation tests to provide a comprehensive whole-scalp assessment while minimizing spatial selection bias and controlling Family-Wise Error Rate (FWER) (Ayrolles et al., 2021; Dumas et al., 2010; Maris & Oostenveld, 2007; Zimmermann et al., 2024). Paired t-statistics were calculated from Fisher-transformed values at each homologous electrode pair–frequency feature. Features exceeding the two-sided cluster-forming threshold of p < .05 were grouped into clusters based on BioSemi 64-channel sensor and frequency adjacency. Cluster mass was defined as the sum of the absolute t-statistics within each cluster. Condition labels were randomly swapped within dyads in 5,000 permutations, with paired t-statistics and clusters obtained for each permutation.

Dyads' inter-brain relative phase values were analyzed using paired Hotelling’s T² statistics (Hotelling, 1951), with cluster mass defined as the sum of the constituent T² values. For both measures, the largest cluster mass from each permutation formed the null distribution. An observed cluster was considered significant when its mass exceeded the 95th percentile of this distribution (p_FWER_ < .05), thereby controlling multiple comparisons across all electrodes and frequency bands within each contrast. Multiple comparisons were additionally corrected across the three condition contrasts using the Benjamini–Hochberg (FDR) correction to address the multiple comparison issue.

**SI.3 Inter-brain PLV surrogate analysis**

To examine whether inter-brain PLV reflected unique dyadic interaction, we compared the observed PLV between real partners with a pseudo-dyadic baseline. We generated 5,000 surrogate datasets by randomly pairing Participant 1 from each dyad with Participant 2 from another dyad within the same respiratory condition. In each surrogate iteration, Participant 2 assignments were randomized without replacement, with original partners excluded. Surrogate PLV was calculated separately for each trial and then averaged across electrodes and frequency bands. One-sided p values were calculated as the proportion of surrogate PLV values equal to or greater than the observed PLV. The mean observed and surrogate PLVs were 0.226 and 0.219, respectively, during normal breathing; 0.229 and 0.220 during in-phase breathing; and 0.229 and 0.220 during anti-phase breathing (all surrogate-test ps < .001, SI. Figure 1).

To examine the contribution of the original temporal alignment between partners, we additionally generated 5,000 within-trial, epoch-shuffled surrogate datasets. This procedure preserved the original partner pairing and trial context while randomly permuting Participant 2’s epochs relative to Participant 1’s epochs without replacement. Surrogate PLV was calculated and averaged as described above. The mean observed and surrogate PLVs were 0.226 and 0.219, respectively, during normal breathing; 0.229 and 0.221 during in-phase breathing; and 0.229 and 0.221 during anti-phase breathing (all surrogate-test p's < .001, SI. Figure 1).

**SI. Figure 1**

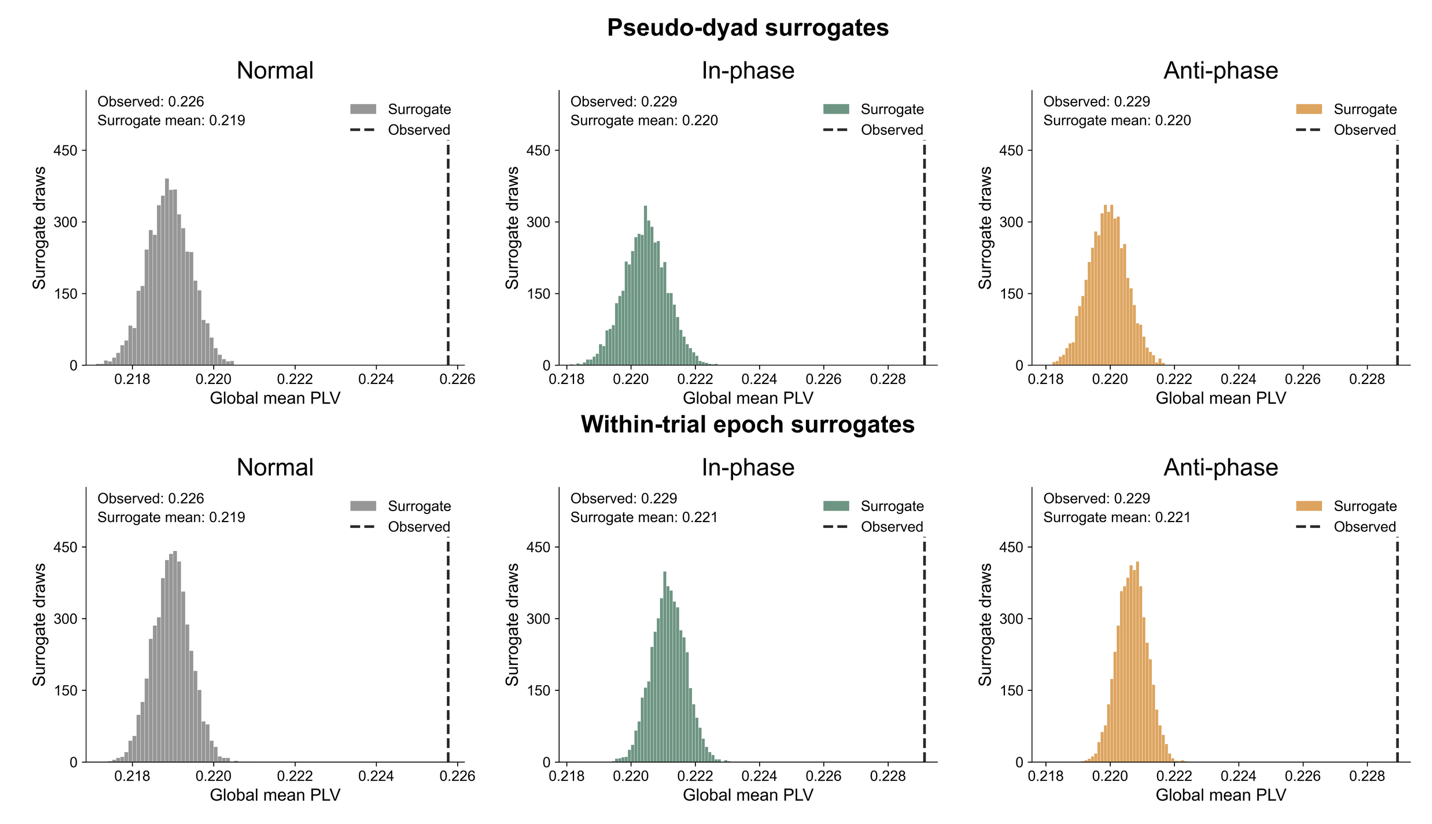

**SI.4 Auditory perceptual screening**

**Method Description:**

Before commencing the experiment, participants completed a Perception Task designed to test whether participants could consistently detect asynchronies in tone onsets. Participants heard 12 trials containing two piano tones separated by one octave. On 1/3 of trials, the two tones began simultaneously (0 ms apart) and on 2/3 of trials, the two tones began asynchronously (80 ms apart).  Tones always ended at the same time, 300 ms after the onset of the first tone.  Participants had to indicate whether the two tones in each trial occurred simultaneously or not. Two practice trials preceded the perception task. Participants were excluded from the study if they answered fewer than 9/12 trials correctly; no participants were excluded in the task.

**SI.5 Cardiac pre-processing and synchronization**

Cardiac synchrony was quantified as the phase-locking value (PLV) between partners’ heart-rate time series. Bipolar ECG signals were resampled to 500 Hz and processed using NeuroKit2 Python for polarity correction, cleaning, and R-peak detection within contiguous segments retained after manual quality control. RR intervals outside 300–2,000 ms (physiologically impossible range for cardiac activity) were rejected, with isolated interior invalid intervals interpolated; segments were excluded when invalid intervals exceeded 20% of their total RR duration. RR intervals were resampled to heart rate BPM (beats/min) and averaged in 3-s windows advanced in 0.25-s steps, yielding equally sampled, equal-length time series for the two partners, and then z-scored within trial; Hilbert-derived phase differences were used to calculate PLV. One dyad was excluded because of noisy ECG recording, leaving 43 complete dyads for repeated-measures ANOVA. Cardiac synchrony differed across conditions (SI. Figure 2), F(2, 84) = 5.28, *p* = .00694, η²_G = .074. Paired comparisons corrected using the Benjamini–Hochberg procedure across the three condition contrasts, showed higher PLV in In-phase and Anti-phase than in Normal respiratory conditions (both *p* = .0111), with no significant difference between In-phase and Anti-phase (*p* = .945). Indicating synchronized respiration rate increases heart rate synchronization.

**SI. Figure 2**

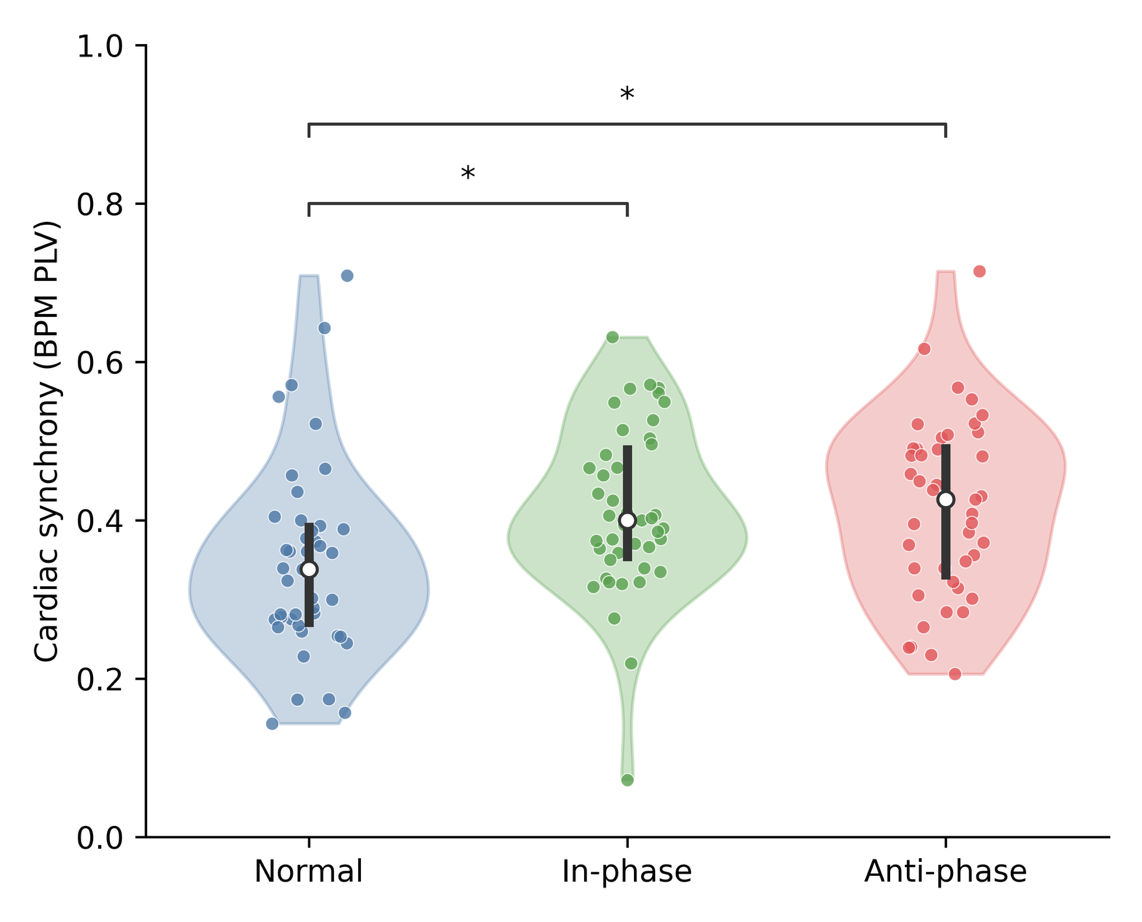

**SI.6 Spontaneous Production Rate Measurement**

Participants produced the melody "Mary Had a Little Lamb" by tapping the force-sensitive pad with their index finger at a steady and consistent rate, with each tap producing the next tone in the sequence at a fixed loudness. Participants were instructed to produce the melody continuously for three iterations per trial, and they completed three trials. The first 13 notes of the first iteration and the final 13 notes of the third iteration were excluded, leaving a central segment equivalent to two complete iterations, in order to capture maximally stable spontaneous rhythmic behavior (Van Kerrebroeck et al., 2025; Yi et al., 2025; Yi & Palmer, 2026; Zamm et al., 2016), The mean and standard error of the intertap intervals (ITIs) were computed to obtain each participant’s SPR, with lower mean ITI indicating a faster rate and higher mean ITI indicating a slower rate.

**SI.7 Respiration Timeseries Pre-processing**

Respiratory data were processed in Neurokit 2 Python. Visually examined respiration time series were filtered using a fourth-order zero-phase Butterworth band-pass filter (0.05–1.0 Hz) and standardized to remove high-frequency noise and low-frequency drift for subsequent analysis, following the established pipeline (Konvalinka et al., 2023; Yi & Palmer, 2026). Respiration rate was estimated using NeuroKit2’s “rsp_process” function with the BioSPPy method.

Extraction of the inhale-exhale cycles was based on identification of the respiratory peaks and troughs the filtered standardized signals using NeuroKit2’s “rsp_findpeaks” function with the SciPy method (minimum peak distance = 0.6 s; prominence = 0.10). Inhalation extended from a peak (local maximum) to the following trough (local minimum), and exhalation extended from that same trough to the next peak. For associated behavioral analyses, the partners’ corresponding respiratory onset and offset times were averaged across the inhale periods and across the exhale periods to define each half-cycle window (Inhale or Exhale). The earliest matched tapping event within ±100 ms of the window onset was included based on respiratory instructions. The analysis included 2,217 inhalation and 2,217 exhalation half-cycles across 44 dyads, containing 6,996 (inhale period) and 7,655 (exhale period) tapping events, respectively. The mean number of tapping events per half-cycle was 3.2 during inhalation and 3.4 during exhalation.
